# Twist-bend coupling and the statistical mechanics of DNA: perturbation theory and beyond

**DOI:** 10.1101/422683

**Authors:** Stefanos K. Nomidis, Enrico Skoruppa, Enrico Carlon, John F. Marko

## Abstract

The simplest model of DNA mechanics describes the double helix as a continuous rod with twist and bend elasticity. Recent work has discussed the relevance of a little-studied coupling *G* between twisting and bending, known to arise from the groove asymmetry of the DNA double helix. Here, the effect of *G* on the statistical mechanics of long DNA molecules subject to applied forces and torques is investigated. We present a perturbative calculation of the effective torsional stiffness *C*_eff_ for small twist-bend coupling. We find that the “bare” *G* is “screened” by thermal fluctuations, in the sense that the low-force, long-molecule effective free energy is that of a model with *G* = 0, but with long-wavelength bending and twisting rigidities that are shifted by *G*-dependent amounts. Using results for torsional and bending rigidities for freely-fluctuating DNA, we show how our perturbative results can be extended to a nonperturbative regime. These results are in excellent agreement with numerical calculations for Monte Carlo “triad” and molecular dynamics “oxDNA” models, characterized by different degrees of coarse-graining, validating the perturbative and non-perturbative analyses. While our theory is in generally-good quantitative agreement with experiment, the predicted torsional stiffness does systematically deviate from experimental data, suggesting that there are as-yet-uncharacterized aspects of DNA twisting-stretching mechanics relevant to low-force, long-molecule mechanical response, which are not captured by widely-used coarse-grained models.

## I. INTRODUCTION

In vivo, double-stranded DNA is typically found in a highly-deformed state, which is in part due to the interaction with the many proteins that bend and twist the double helix, but in part due to thermally-driven deformations. A substantial effort has been devoted to the study of many aspects of DNA mechanics, such as its response to applied twist and bending deformations [1]. These studies often rely on homogeneous elastic models, which, despite their simplicity, describe many aspects of single-molecule experiments [2–6], and are widely used to describe mechanical and statistical-mechanical properties of DNA (see e.g. Refs. [7–11]).

One of the simplest models describing DNA deformations is the twistable wormlike chain (TWLC), which describes the double helix as an inextensible rod, for which twist and bend deformations are independent. Symmetry arguments suggest that the TWLC is incomplete: The inherent asymmetry of the DNA molecular structure, with its major and minor grooves, gives rise to a coupling *G* between twisting and bending [12]. Only a limited number of studies have considered the effect of twist-bend coupling on DNA mechanics [13–16]. A systematic analysis of coarse-grained models with and without groove asymmetry have highlighted several effects associated with twist-bend coupling at long [15] and short [16] length scales. Here, we aim to clarify the role of *G* in the statistical mechanics of long DNA molecules, as analyzed in optical and magnetic tweezers.

We focus on analytical and numerical results for the stretching and torsional response of DNA with twist-bend coupling interaction *G* ≠ 0. We first present a perturbative expansion for the partition function of the molecule, in which *G* is treated as the small parameter. The lowest-order results show that twist-bend coupling softens the torsional and bending stiffnesses of the double helix, recovering prior results from entirely different calculations [14]. Our new calculations reveal the existence of a previously-unidentified large force scale *f*_0_; for forces below this scale, the bare elastic constants - including *G* - are not directly accessible in stretching and twisting experiments. Instead, for forces below *f*_0_, only renormalized bending and twisting stiffnesses - which do depend on *G* - are observed. Because *f*_0_ ≈ 600 pN, the renormalized elastic model - which is the *G* = 0 TWLC - will be observed in essentially all conceivable single-molecule experiments. Thus, *G* is “screened”, effectively renormalized to *G* = 0, in single-molecule DNA mechanics experiments.

Prior work [14] suggests a strategy to generalize our results beyond perturbation theory, to the regime where DNA is stretched by forces less than *f*_0_. We validate both the perturbative and nonperturbative results using numerical calculations corresponding to commonly-used coarse-grained DNA elasticity models; our results turn out to closely describe results of those numerical models. Given this validation, we turn to experimental data which are reasonably well described by the low-force model, but for which there remain discrepancies, suggesting effects beyond simple harmonic elastic models like the TWLC.

## II. ELASTICITY MODELS OF DNA

To describe the conformation of a continuous, inextensible, twistable elastic rod, one can associate a local orthonormal frame of three unit vectors {**ê***_i_*} (*i* = 1,2,3) with every point along the rod (Fig. 1). In a continuous representation of DNA, the common convention is to choose **ê**_3_ tangent to the curve **ê**_1_ and pointing to the DNA groove. The frame is completed with a third vector, defined as **ê**_2_ = **ê**_3_ × **ê**_1_. An unstressed B-form DNA corresponds to a straight, twisted rod, with the tangent **ê**_3_ being constant, and with **ê**_1_ and **ê**_2_ rotating uniformly about it, with a full helical turn every *l* ≈ 3.6 nm, or equivalently every 10.5 base pairs.

**FIG. 1.**
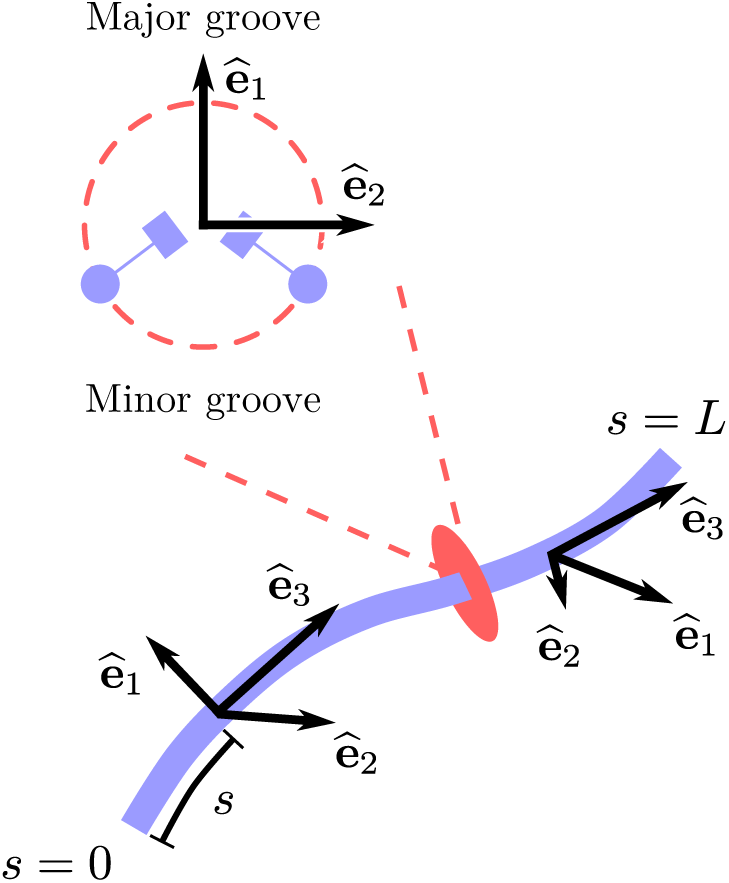
Bottom: The configuration of a twistable elastic rod can be mathematically described with an orthonormal set of vectors {**ê**_1_, **ê**_2_, **ê**_3_} assigned to every point *s* along the rod. The vector **ê**_3_ is the tangent to the curve, and describes the bending fluctuations along the rod. Top: Cross-section of the rod, indicating how the remaining vectors **ê**_1_ and **ê**_2_, which describe the torsional state, may be chosen in the particular case of DNA.

Any deformation from this unstressed configuration can be described by a continuous set of rotation vectors **Ω** connecting adjacent local frames {**ê***_i_*} along the rod, using the differential equation

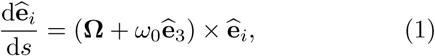

where the internal parameter *s* denotes the arc-length coordinate (Fig. 1), and *ω*_0_ = 2*π/l* ≈ 1.75 nm^−1^ is the intrinsic twist of DNA. Upon setting **Ω** = **0**, one obtains the unstressed configuration mentioned above. Thus, a nonzero rotation vector **Ω**(*s*) ≠ **0** corresponds to a local deformation at *s* around this ground state. Defining Ω*_i_* ≡ **ê***_i_*. **Ω**, it follows that Ω_1_(*s*) and Ω_2_(*s*) describe local bending deformations, while Ω_3_(*s*) describes twist deformations. In the remainder of the paper the *s*-dependence of **Ω** will be implicit.

Symmetry analysis of the DNA molecule requires the energy functional *E* to be invariant under the transformation Ω_1_ →-Ω_1_, with the consequence that [12]

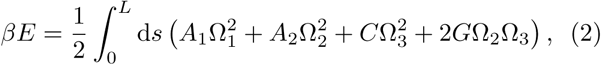

where *β* ≡ 1*/k*_B_*T* is the inverse temperature, *A*_1_ and *A*_2_ the bending stiffnesses, *C* the torsional stiffness and *G* the twist-bend coupling constant. These coefficients have dimensions of length, and can be interpreted as the contour distance along the double helix over which significant bending and twisting distortions can occur by thermal fluctuations. Our perturbative calculation will use the isotropic-bending version of this model (*A*_1_ = *A*_2_ = *A*), which is described by the following energy functional

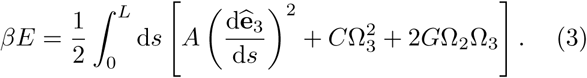

Here, we have used Eq. (1) to express the sum 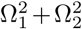 as the derivative of the tangent vector. The TWLC is obtained by setting *G* = 0 in Eqs. (2) and (3), corresponding to the anisotropic and isotropic cases, respectively.

## III. EFFECTIVE TORSIONAL STIFFNESS

In a typical magnetic tweezers experiment, a single DNA molecule of 10^3^ – 10^4^ bases is attached to a solid substrate and to a paramagnetic bead at its two ends (Fig. 2). The molecule can be stretched by a linear force *f* and over- or undertwisted by an angle *θ*. The resulting torque *τ* exerted by the bead, which can be experimentally measured [17–20], is linear in *θ* for small *θ*

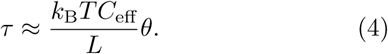

**FIG. 2.**
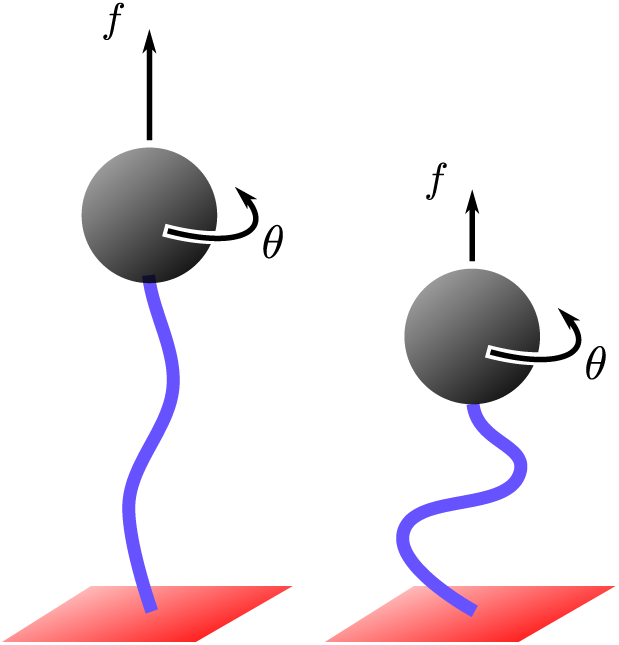
Typical setup of a magnetic tweezers experiment. A DNA molecule is covalently bound to a substrate at one end and to a paramagnetic bead at the other end. An applied force f stretches the molecule, while an applied rotation θ twists it. The effective torsional stiffness *C*_eff_ is the proportionality constant connecting the applied rotation with the exerted torque [Eq. (4)].

Here *C*_eff_ is the *effective* torsional stiffness (in contrast to the intrinsic stiffness C), and represents the central quantity of interest here. It expresses the resistance of the DNA to a global torsional deformation, applied at its two ends.

As discussed in more detail below, *C*_eff_ is in general lower than its intrinsic equivalent *C*. More specifically, at low stretching forces the bending fluctuations can absorb a significant part of the applied torsional stress, leading to a globally-reduced torsional resistance *C*_eff_ < *C*. On the other hand, when the applied force is sufficiently large, bending fluctuations are mostly suppressed, and hence the effective torsional stiffness tends to approach the intrinsic one. As a consequence, *C*_eff_ is going to be a monotonically-increasing function of the stretching force.

Moroz and Nelson derived an expression of *C*_eff_ for the TWLC in the limit of high forces [4, 21]. In spite of the good qualitative agreement between the theory and early experiments, more recent studies reported systematic deviations [14, 17, 20, 22]. For completeness we will first present in Sec. III A a short derivation of the TWLC-based theory by Moroz and Nelson. The pertubative calculation in small *G* is discussed in Sec. III B and generalized beyond perturbative expansion in Sec. III D.

### A. The TWLC limit (*G* = 0)

In their original calculation, Moroz and Nelson [4, 21] mapped the twisted and stretched TWLC onto a quantum mechanical problem of a spinning top, and *C*_eff_ was obtained from the ground state of the associated Schrödinger equation. Here we present an alternative derivation, following the scheme illustrated in Ref. [1], which proves to be more convenient for the perturbative calculation in small *G*. The starting point is the partition function of a TWLC under applied force *f* and torque *τ*. The latter induces a rotation by an angle *θ* on the end point of the molecule (Fig. 2). The excess linking number, which we will use throughout this work, is ΔLk = *θ*/2*π*.

To calculate the partition function, we integrate over all possible configurations of the twistable rod, which can be parametrized by the tangent vector **ê**_3_ (*s*) and the twist density Ω_3_(*s*). The resulting path integral takes the form:

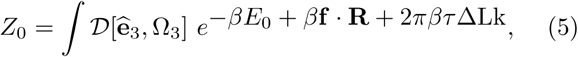

where *E*_0_ is the energy of the TWLC, obtained from Eq. (3) by setting *G* = 0, and **R** the end-to-end vector

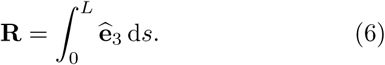

we assume that the force is oriented along the z-direction, hence 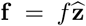, with 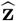 a unit vector. Using a result due to Fuller [23] for open curves, the linking number can be expressed as the sum of twist and writhe, i.e. ΔLk = Tw + Wr. The excess twist is obtained by integrating over the twist density

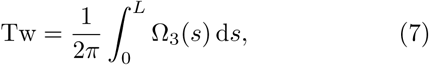

while the writhe is given by [1]

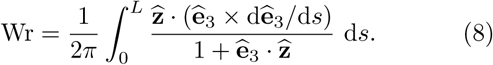

This representation of the writhe as a single integral (and not as a double, nonlocal integral, as in closed curves) is correct modulo 2*π*, and possible if the molecule is sufficiently stretched and does not loop back opposite to the direction of the applied force. Under these conditions, the denominator 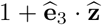 does not vanish, and the integral in Eq. (8) yields a finite value. Next, we insert Eqs. (6), (7) and (8) into Eq. (5), and consider the limit of strong forces and weak torques. The partition function [Eq. (5)] reduces to a Gaussian in this limit, and can be easily estimated (details can be found in Appendix A). To lowest order in *τ* and at large forces, one obtains the following free energy

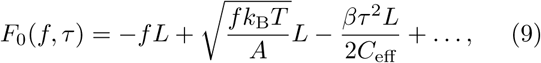

where the dots denote constant or higher-order terms in *τ*. The effective torsional stiffness *C*_eff_ is given by (see Appendix A)

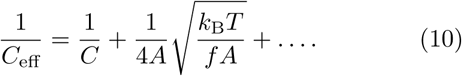

This equation was originally derived by Moroz and Nelson [4, 21]. At high forces, Eq. (10) approaches the twisted-rod limit and *C*_eff_ → *C*, but, in general, bending fluctuations soften the DNA torsional stiffness, so that *C*_eff_ <*C*. The latter originates from a global coupling between torque and writhe (not to be confused with the local twist-bend coupling considered below). Note that the effect of bending fluctuations is governed by the dimensionless parameter 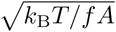, which is small at low temperatures or high forces.

### B. Perturbative (small-*G*) expansion

We now construct a perturbation expansion for the partition function, using *G* as the small parameter (the length scale determining whether *G* is “small” will be made clear below). The full partition function is

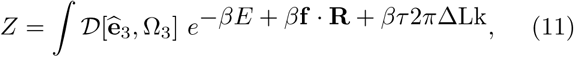

where now *βE* is given by Eq. (3) and contains a twist-bend coupling term. Assuming that *G* is small, we can expand the Boltzmann factor in powers of *G*, which gives to lowest order:

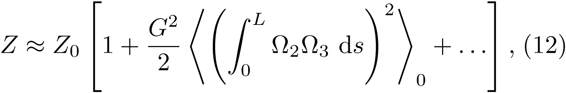

where ⟨.⟩_0_ denotes the average with respect to the unperturbed (TWLC) partition function [Eq. (5)]. Note that in the perturbative expansion, the term linear in *G* vanishes by the Ω_2_ → –Ω_2_ symmetry of the TWLC. The full calculation of the average in the right-hand side of Eq. (12) is given in Appendix C. The final expression for the free energy is of the form [Eq. (C38)]

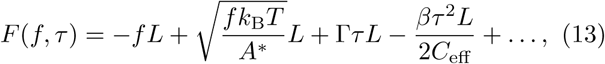

where terms of negligible contribution were omitted (see Appendix C).

In the above, we have introduced the rescaled bending stiffness

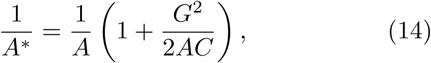

together with the parameter

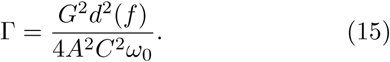

The dimensionless, force-dependent scale factor *d*(*f*) will be discussed below; we note that it appears in the coefficient Г [Eq. (15)], but not in *A*^*^ [Eq. (14)]. Finally, the form of the effective torsional stiffness *C*_eff_ is

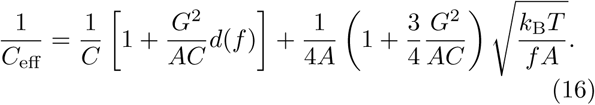

The scale factor *d*(*f*) is also present in *C*_eff_.

Examination of these formulae indicate that the expansion is in powers of the dimensionless parameter *G*^2^*/AC*, which, given our current estimates for the stiffnesses (*A* ≈ 50 nm, *C* ≈ 100 nm, *G* ≈ 30 nm), is less than 1, although we note that *G*^2^*/AC <* 1 is a stability requirement for the microscopic energy [12]. Our computation neglects terms beyond first order in *G*^2^*/AC*.

### C. Effective torsional stiffness *C*_eff_

Equation (16) is the central result of this paper, and extends the TWLC result by Moroz and Nelson [Eq. (10)], which is recovered in the the limit *G* → 0. The perturbative corrections are governed by the dimensionless parameter *G*^2^*/AC*, and give rise to a further torsional softening of the molecule, i.e. *C*_eff_(*G* ≠ 0) <*C*_eff_(*G* = 0), as pointed out in Ref. [14]. Equation (16) contains also a force-dependent, crossover function, which can be approximated as (see Appendix C)

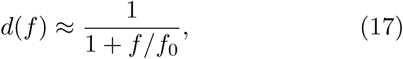

where 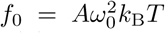 is the characteristic force above which *d*(*f*) starts to significantly drop below its low-force limit of *d*(0) = 1. To understand this force scale, which has no counterpart in the Moroz and Nelson formula [Eq. (10)], we recall that the correlation length for a stretched wormlike chain is 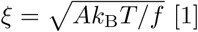 [1]. Therefore, *f*_0_ is the force associated with a correlation length of the order of the distance between neighboring bases, i.e. *ξ* = 1*/ω*_0_.

For DNA (*A* ≈ ≈ 50 nm, *ω*_0_ = 1.75 nm^−1^, *k*_B_*T* = 4 pN·nm) we see *f*_0_ ≈ 600 pN, which is far above the force where the double helix starts to be itself stretched (≈ 20 pN), force-denatured (≈60 pN), and is in fact comparable to where the covalently-bonded backbones will break. Hence, for forces relevant to experiments we are concerned with (*f* < ≈ 10 pN), one may simply set *d*(*f*) ≈ 1. We will refer to this limit as the “low-force limit”, but one should keep in mind that our perturbative theory is computed for the “well-stretched” limit, i.e. *f* >*k*_B_*T/A* ≈ 0.1 pN. Therefore our perturbative theory is applicable in the force range of roughly 0.1 to 10 pN.

We emphasize that Eq. (16) can be written as

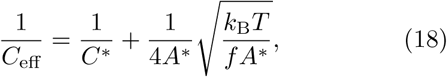

where *A*^*^ is defined in (14), and

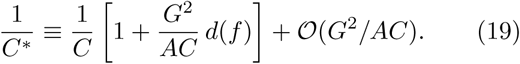

Equation (18) has exactly the same form as the Moroz and Nelson formula [Eq. (10)], with rescaled bending and torsional stiffnesses. The importance of this result is paramount: since *d*(*f*) = 1 in the range of experimentally relevant forces, the torsional stiffness (and in fact the partition function itself) depends only on the “renormalized” stiffnesses *A*^*^ and *C*^*^, meaning that *G* by itself cannot be determined from fitting of *C*_eff_(*f*) (or any other equilibrium quantity versus *f*); only the effective stiffnesses *A*^*^ and *C*^*^ can be determined from experiments in the low-force regime.

### D. Non-perturbative result for *C*_eff_ valid for *f* <*f*_0_

Equation (16) has been derived on the basis of a systematic perturbation expansion in *G* (more formally, in the small parameter *G*^2^*/AC* < 1). When cast in the form of Eq. (18), it is apparent that there is a simple way to extend the results to a more general, nonperturbative case, where *G* may be large and the bending possibly anisotropic, i.e. *A*_1_ ≠ *A*_2_ in Eq. (2). The key physical idea here is that for forces below the gigantic force scale *f*_0_, thermal fluctuations at the helix scale where *G* correlates bending and twisting fluctuations is unperturbed (*d*(*f*) = 1), and therefore we might as well just consider DNA to have the effective twisting and bending stiffnesses that is has at *zero* force.

In absence of applied torques and forces (*f, τ* = 0), the partition function of Eq. (11) can be evaluated exactly [14]. Due to twist-bend coupling, the bending and torsional stiffnesses are renormalized as [14]:

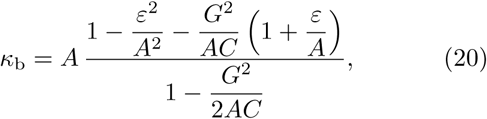

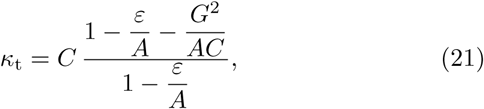

where we have introduced the parameters *A* = (*A*_1_ + *A*_2_)/2 and *ε* = (*A*_1_ − *A*_2_)/2. Eqs. (20) and (21) quantify the energetic cost of bending and twisting deformations, respectively, in the same way *A* and *C* do within the TWLC. Note that, by setting *G* = 0 in these expressions, one recovers the TWLC limit, *κ*_t_ = *C* and *κ*_b_ = 2*A*_1_*A*_2_/(*A*_1_ + *A*_2_) i.e. the renormalized bending stiffness is the harmonic mean of *A*_1_ and *A*_2_, which is a known result (see e.g. Refs. [24, 25]). If *G* ≠ 0 one has *κ*_b_ < 2*A*_1_*A*_2_/(*A*_1_ + *A*_2_) and *κ*_t_ <*C*, i.e. twist-bend coupling “softens” the bending and twist deformations of the DNA molecule, already if *f* = 0, *τ* = 0. Eq. (16) then describes two different effects: one is the thermally-induced torsional softening due to bending fluctuations [already present in the TWLC expression (10)] and the other is the *G*-induced softening, which is captured by the two factors between parentheses in Eq. (16).

Setting *ε* = 0 and expanding Eqs. (20) and (21), onefinds *κ*_b_ = *A*^*^ + 𝒪(*G*^4^) and *κ*_t_ = *C*^*^ + 𝒪(*G*^4^), which suggests the following, more general, nonperturbative result for *C*_eff_, valid as long as *f* ≪ *f*_0_

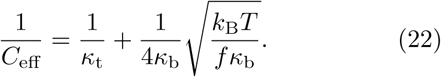

This relation, similar to Eq. (18), has the same form as the Moroz and Nelson formula [Eq. (10)], with *A* and *C* replaced by *κ*_b_ and *κ*_t_ [much as our result for *C*_eff_ of Eq. (19) has the Moroz-Nelson form with *A* → *A*^*^ and *C* → *C*^*^]. As we will show in the next Section, this new, nonperturbative result for the continuum model is in excellent agreement with numerical Monte Carlo (MC) and molecular dynamics (MD) calculations.

### E. Twist-bend-coupling-induced DNA unwinding

An intriguing feature of the perturbative calculation is the appearance of a term linear in *τ* in the free energy [Eq. (13)], which induces an unwinding of the helix at zero torque. In particular, from Eqs. (11), (13) and (15) it follows that

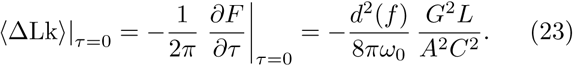

The scale for this thermal unwinding is very small: using typical values of DNA parameters (*A* ≈ 50 nm, *C* ≈ 100 nm, *G* = 30 nm, *ω*_0_ = 1.75 nm ^−1^) we find an unwinding angle per contour length of 2*π* Δ k */L* = *G*^2^/4*ω*_0_*A*^2^*C*^2^ 5 × 10^−6^ rad/nm (about 1 10^−5^ degrees per base pair). This *G*-generated shift in helix twisting is inconsequentially small, but it is worth noting that this term is present in the perturbation theory.

It has been long known that there is a gradual unwinding of the double helix as the temperature is increased [26, 27], and this effect has been recently observed at the single-DNA level [28]. Although one might imagine an overall *T* ^2^ dependence of this term (from the factors of *β* in the Boltzmann factor), this dependence can only generate a tiny fraction of the observed temperature-dependent unwinding of ≈ –1 × 10^−2^ degrees/K·bp. The experimentally observed unwinding is likely due to temperature-dependence DNA conformational changes [28], and is beyond the scope of being captured by the simple elastic models discussed here; in particular the observed unwinding of DNA with increasing temperature is not attributable to the twist-bend coupling *G*.

### F. “Janus strip” limit (*ω*_0_ → 0)

Equation (22) is not generally valid for any arbitrary polymer with twist-bend coupling, but its validity is linked to the physical parameters characterizing DNA elasticity. These conspire to set forces encountered in typical experiments (below 10 pN) to be far below the characteristic force 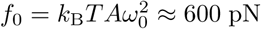 pN at which one starts to see effects at the helix repeat scale, i.e. force-driven unwinding of the double helix due to quenching of thermal fluctuations and the influence of *G*. In this sense, *ω*_0_ can be regarded as a “large” parameter: combinations of it and the elastic constants give dimensionless constants large compared to unity, e.g. *Aω*_0_, *Cω*_0_ ≈ 10^2^ ≫ 1 and 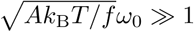 for *f* <10 pN).

For this reason, several *ω*_0_-dependent terms, which in principle would contribute to *C*_eff_ at order *G*^2^, can in practice be neglected in the application of the theory to DNA [see e.g. Eqs. (C30) and (C31) in Appendix]. The neglect of these terms leads to *C*_eff_ taking the simple form given by Eq. (18), in which *A*^*^ and *C*^*^ are the renormalized stiffnesses.

While not relevant to DNA, we might imagine other polymer structures for which *ω*_0_ is not so large, i.e. where *ω*_0_ is closer in size to 1*/A* or 1*/C*. In this case one cannot ignore these additional terms, and *d*(*f*) ≈ 1/(1+*f*/*f*_0_) might drop significantly over experimentally-relevant force ranges. Chiral proteins, lipid filaments, or even nanofabricated objects might comprise realizations of such situations.

As an example we consider the extreme limit *ω*_0_ → 0, corresponding to a “Janus strip”, an elastic strip with inequivalent faces (i.e. inequivalent major and minor “grooves”), and, thus, nonzero *G*. In this case, using the more complete and complicated results for the perturbative expansion given in the Appendix, we obtain to lowest order in *G*

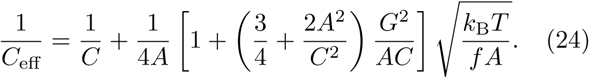

In this limit, compared to the large-*ω*_0_ case relevant to DNA, there is a more gradual shift of *C*_eff_ up to its high-force limit, and an inequivalence of the form of *C*_eff_ to the Moroz-Nelson form. Physically, this is because the intrinsic chirality of the filament is now gone, eliminating the “screening” of effects of *G* at low forces, and the simple dependence of the low-force themodynamics on only the coarse-grained stiffnesses *κ*_t_ and *κ*_b_. Experiments on such Janus strips, or on “soft-helix” objects where *Aω*_0_, *Cω*_0_ *<* 1 could provide realizations of this limit of the theory. Ref. [14] showed that Eq. (24) fits experimental data for DNA surprisingly well, despite not taking account of double helix chirality.

## IV. NUMERICAL CALCULATIONS

To check the validity of the analytical results presented above, we performed numerical simulations of two different models. The first model, referred to as the *triad model*, is obtained from the discretization of the continuum elastic energy (2) and treated using MC computations. The second model is oxDNA, a coarse-grained model of nucleic acids [29], treated using MD calculations.

### A. Triad model

The triad model is comprised of a series of *N* orthonormal vectors {****ê*****_i_*} with *i* = 1, 2, 3 and *k* = 0, 1, 2 *… N*, each representing a single base pair, interacting with its neighbors according to Eq. (2). The total length of the molecule is *L* = *Na*, with *a* = 0.34 nm the base pair distance. The ground state of this model is a twisted, straight rod, with ****ê****_3_ being aligned with the direction of the stretching force, and the vectors ****ê****_1_, ****ê****_2_ rotating about ****ê****_3_ with an angular frequency *ω*_0_. A cluster move consisted of a rotation of the whole subsystem beyond a randomly-selected triad by a random angle. The new rotation vector **Ω** was calculated based on Rodrigues’ rotation formula (see e.g. Supplementary Material of Ref. [15]), then the energy was updated from a discretized version of Eq. (2), with the addition of a force term [see Eq. (5)]. The move was accepted or rejected according to the Metropolis algorithm. The stiffness constants *A*_1_, *A*_2_, *C* and *G* are input parameters for the model, and may, therefore, be arbitrarily chosen, provided the stability condition *G*^2^ <*A*_2_*C* is met [for which the quadratic form in Eq. (2) is positive definite]

The effective torsional stiffness was calculated at zero torque from linking number fluctuations:

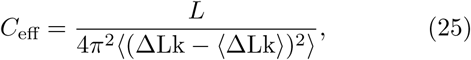

The variance of linking number in the denominator was evaluated from the topological relation ΔLk = ΔTw + Wr, with twist and writhe obtained from the discretization of Eqs. (7) and (8), respectively. To check the validity of our results, the writhe was also evaluated from the double-integral formula, following the method of Ref. [30], and no significant differences were found for forces > 0.25 pN. In all simulations, the size of the system was 600 triads (base pairs), above which the results remained identical within that force range.

#### 1. Isotropic bending

Figure 3 shows the results of Monte Carlo calculations for the isotropic model of Eq. (3), with *A* = 50 nm, *C* = 100 nm and *G* = 0, 20, 40 nm (top to bottom). The data are plotted as a function of the dimensionless parameter 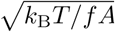. The numerical errors are smaller than the symbol sizes, and hence not shown.

**FIG. 3.**
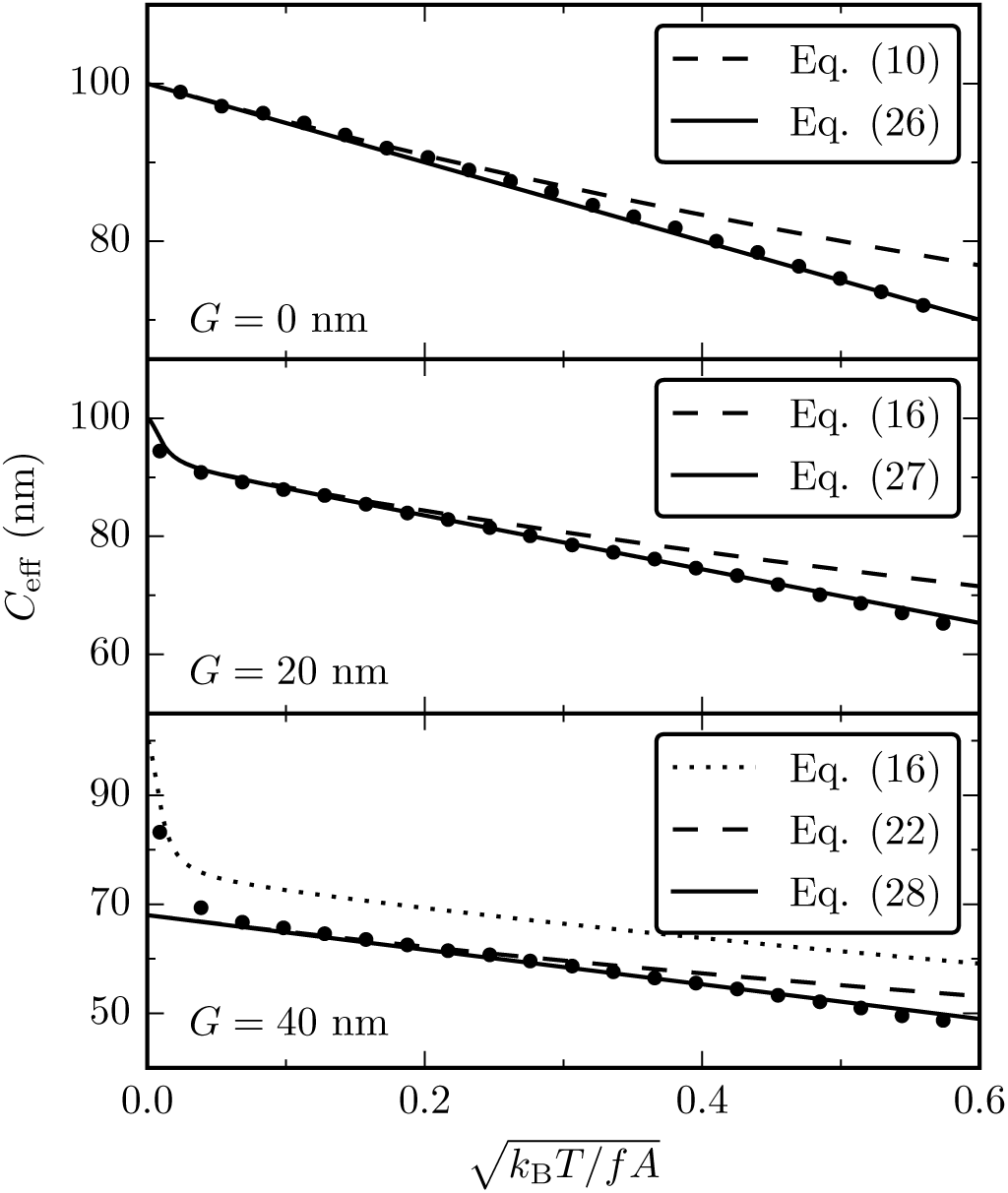
Comparison of Monte Carlo simulations of the triad model (points) with various analytical expressions of *C*_eff_ (lines) for *A* = 50 nm, *C* = 100 nm and *G* = 0, 20, 40 nm. The data are plotted as a function of 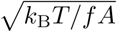 and correspond to *f* ≥ 0.25 pN. The numerical results are in excellent agreement with the analytical expressions, both in the perturbative and nonperturbative regimes (see text). Error bars of Monte Carlo data are smaller than symbol sizes.

In absence of twist-bend coupling (*G* = 0, upper panel), the Monte Carlo data are in excellent agreement with the Moroz-Nelson theory. We compare them both to Eq. (10) (dashed line) and the following expression (solid line)

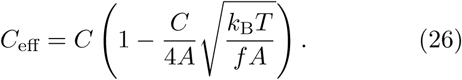

The latter is obtained from the lowest-order expansion of Eq. (10) in 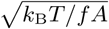, and is a straight line when plotted as a function of the rescaled variable of F<u>ig</u>. 3. Eqs. (10) and (26) coincide to leading order in 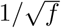, and any differences in the two expressions only occur at low force scales, where higher-order corrections become relevant. Eq. (26) fits the Monte Carlo data extremely well over the whole range of forces analyzed (*f* ≥ 0.25 pN), while Eq. (10) deviates at low forces.

The middle panel of Fig. 3 shows Monte Carlo results for *G* = 20 nm (points), which we compare both to the results for the perturbative expansion for 1*/C*_eff_, Eq. (18) (dashed line) and the following similar expansion result for *C*_eff_ (solid line)

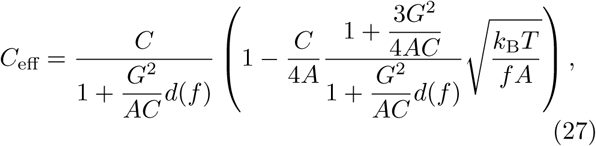

obtained by expanding Eq. (16) to lowest order in 1*/f*. The latter is in excellent agreement with Monte Carlo data in the whole range of forces considered, indicating that *G* = 20 nm falls within the range of validity of the perturbative calculation (*G*^2^*/AC* = 0.08 in this case). We note that the two perturbative expansion results converge together at high forces ([*k*_*B*_*T*/(*Af*)]^1/2^ → 0 and also show the upturn at the very highest forces associated with the force-dependence of *d*(*f*).

Finally, the lower panel of Fig. 3 shows the results of Monte Carlo simulations for *G* = 40 nm. The numerical data deviate substantially from Eq. (16) (dotted line), indicating that *G* is leaving the range of validity of the perturbative calculation (*G*^2^*/AC* = 0.32). The remaining curves show the nonperturbative result for 1*/C*_eff_ Eq. (22) (dashed line), together with the following nonperturbative expression for *C*_eff_ (solid line)

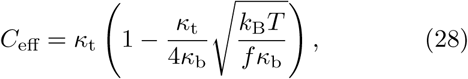

the latter being in excellent agreement with Monte Carlo data for *f* < *f*_0_, validating the nonperturbative result (note that [*k*_*B*_*T*/(*f*_0_*A*)]^1/2^ 0.01). For *f* > *f*_0_, the up-turn of *C*_eff_ towards the bare value of *C* is apparent; this effect, while not given by the nonperturbative results, is present in the perturbation expansion results. We conclude that our nonperturbative result indeed provides a quantitative account of *C*_eff_ for *f < f*_0_, where we expect it to be valid.

#### 2. Anisotropic bending

While the perturbative calculation [Eq. (16)] was restricted to the isotropic case (*A*_1_ = *A*_2_), the nonperturbative result [Eq. (28)] has a broader range of applicability, and is able to describe the anisotropic case as well. Figure 4 shows the results of Monte Carlo calculations of *C*_eff_ for various values of the anisotropy parameter *ε* = (*A*_1_ − *A*_2_)/2 and a fixed value of the force *f* = 1 pN. The Monte Carlo data are in very good agreement with Eq. (28), plotted with solid lines. The differences are within 5%, and are probably due to higher order corrections in 1*/f* (recall that all analytical results are based on a large-force expansion). In absence of twist-bend coupling (*G* = 0), Eq. (28) is symmetric in *ε*, as in this case Eqs. (20) and (21) give *κ*_b_ = (*A*^2^ – *ε*^2^)*/A* and *κ*_t_ = *C*, respectively. A nonzero *G* induces nonvanishing terms, which are linear in *ε* both in *κ*_b_ and *κ*_*t*_ [Eqs. (20) and (21)], leading to a breaking of the *ε* → *-ε* symmetry.

**FIG. 4.**
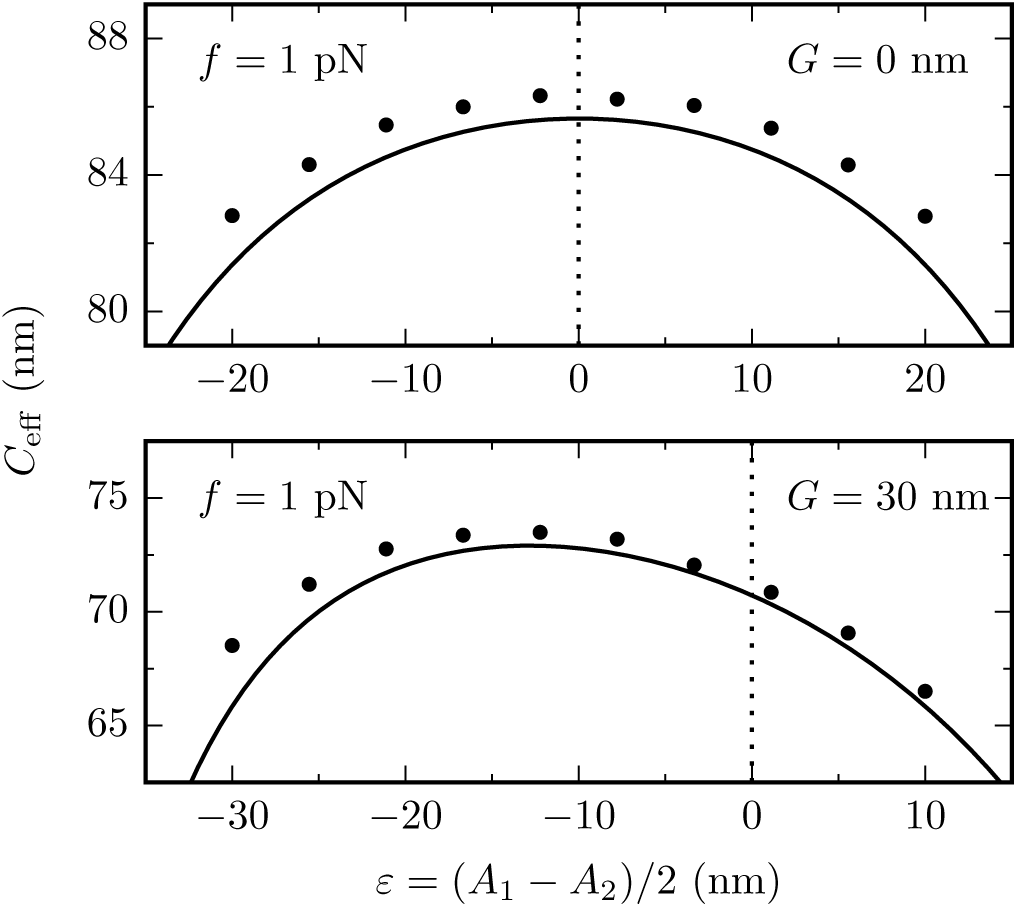
Effect of bending anisotropy (*A*1 ≠ *A*2) on the effective torsional stiffness at a fixed force *f* = 1 pN. Numerical data from Monte Carlo simulations of the triad model (points) are in good agreement with the analytical, nonperturbative predictions of Eq. (28) (solid lines). The vertical dashed lines indicate the isotropic case (*ε* = 0). A nonvanishing twist-bend coupling (lower panel) induces a *ε* →–*ε* symmetry breaking. Error bars of Monte Carlo data are smaller than symbol sizes.

### B. oxDNA

oxDNA is a coarse-grained model describing DNA as two intertwined strings of rigid nucleotides [29]. It has been used for the study of a variety of DNA properties, ranging from single molecules to large-scale complexes [15, 16, 29, 31–33]. To date, two versions of oxDNA exist: one with symmetric grooves (oxDNA1) [29] and one with asymmetric grooves (oxDNA2) [32]. Comparing the torsional response of the two versions will allow us to infer the effect of the groove asymmetry on *C*_eff_. Differently from the triad model, in which the stiffness constants *A*_1_, *A*_2_, *C* and *G* are input parameters, in oxDNA they are determined by the molecular force fields used. These force fields were accurately tuned so that the experimental DNA structural, mechanical and thermodynamic properties (as persistence length, melting temperatures and torque-induced supercoiling) are well reproduced [29]. As for real DNA, for oxDNA the elastic constants are emergent via coarse-graining of fluctuations of smaller-scale, molecular motion degrees of freedom.

The stiffness parameters of oxDNA were recently estimated from the analysis of the equilibrium fluctuations of an unconstrained molecule [15], and are shown in Table I (the values of the elastic constants for oxDNA2 shown are the result of transformation of the values obtained in Ref. [15] for the helical coordinate system used in that paper, to the non-helical coordinate system of this paper; see Appendix D). In line with the symmetry arguments of Ref. [12], twist-bend coupling is absent in oxDNA1 (symmetric grooves), while its magnitude is comparable to that of the elastic constants *A*_1_, *A*_2_ and *C* in oxDNA2 (asymmetric grooves). Table I also reports the values of *κ*_b_ and *κ*_t_, which can be obtained in two different, yet consistent, ways [15]: either indirectly from Eqs. (20) and (21), by plugging in *A*_1_, *A*_2_, *C* and *G* of Table I, or directly from the analysis of the corresponding correlation functions in simulations (*κ*_b_ and 2*κ*_t_ are, respecively, the bending and twist stiffnesses [14]).

**TABLE I.** Values of the stiffness coefficients for oxDNA1 and oxDNA2 (expressed in nm), derived from MD data of Ref. [15]. oxDNA1, which has symmetric grooves, is characterized by a negligible twist-bend coupling constant, while *G* = 25 nm for oxDNA2, which has asymmetric grooves. The values of *κ*_b_ and *κ*_t_ are obtained from Eqs. (20) and (21), respectively, and were found to agree with direct computations of those quantities [15]. Note that the oxDNA2 stiffness coefficients have been transformed to coordinates compatible with this paper, see Appendix D).

| | $A_1$ | $A_2$ | $C$ | $G$ | $\kappa_b$ | $\kappa_t$ |
| --- | --- | --- | --- | --- | --- | --- |
| oxDNA1 | 84(14) | 29(2) | 118(1) | <0.3 | 43 | 118 |
| oxDNA2 | 85(10) | 35(2) | 109(1) | 25(1) | 44 | 92 |

Figure 5 shows a plot of the effective torsional stiffness as a function of the applied force, both for oxDNA1 (circles) and oxDNA2 (triangles). *C*_eff_ was evaluated using twist fluctuations via Eq. (25). At large forces, and in agreement with the experimental evidence, oxDNA undergoes a structural transition, hence the simulations were restricted to *f* ≤ 10 pN. The solid and dotted lines of Fig. 5 are plots of Eq. (28) using *κ*_b_ and *κ*_t_ from Table I. For oxDNA1 there is an excellent agreement between the nonperturbative theory and simulations. In this case the nonperturbative theory reduces to the Moroz-Nelson result, with *κ*_t_ = *C* and *κ*_b_ = *A*(1–*ε*^2^*/A*^2^); the good account of oxDNA1 *C*_eff_ by this formula was noted previously (see Ref. [34], Fig. S7). We note that in the light of our present results, this good agreement validates the use of the values of the stiffness parameters obtained in Ref. [15].

**FIG. 5.**
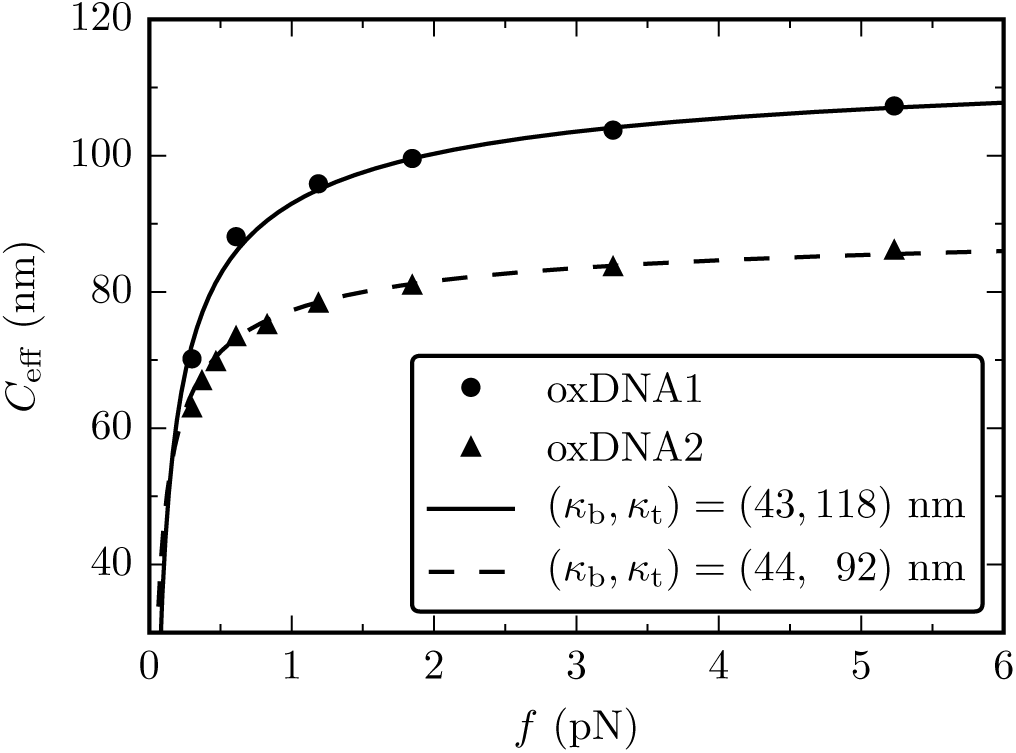
Comparison of oxDNA simulations to Eq. (28). Solid and dashed lines show the nonperturbative result for *C*_eff_ for oxDNA1 and oxDNA2 values of Table I, respectively. Error bars for the oxDNA data are smaller than symbol sizes.

## V. DISCUSSION

We have investigated the effect of the twist-bend coupling *G* on the statistical-mechanical properties of a stretched DNA molecule, using analytical and numerical methods. Our major analytical results are based on a perturbative calculation of the effective torsional stiffness *C*_eff_, the torsional resistance of a long DNA molecule stretched by an applied force *f*. The calculation is valid for small values of *G*, and generalizes the expression derived by Moroz and Nelson, which was derived for *G* = 0 [4].

### A. Screening effect for *f* < *f*0

A striking feature of our theory is the appearance of a large force scale, *f*_0_ = *k*_*B*_*TAω*^2^ ≈ 600 pN. For forces well below this gargantuan force level (essentially all single-DNA mechanical experiments concern forces far below this value) the effect of *G* becomes solely renormalization of the bending and twisting stiffnesses *κ*_b_ and *κ*_t_; direct effects of twist-bend coupling are “screened” at lower force scales. Only at forces *f* ≫ *f*_0_ do the bare elastic constants start to reveal themselves: in this regime *C*_eff_ finally approaches its intrinsic value *C*.

This “screening” feature of the perturbative theory suggested to us that we could consider DNA for *f* < *f*_0_ to be described by a TWLC with persistence lengths set to the zero-force long-molecule stiffnesses *κ*_b_ and *κ*_t_. Combining formulae for the stiffnesses for freely fluctuating DNA [14] with the *C*_eff_ formula of Moroz and Nelson [4] gave us a nonperturbative formula for *C*_eff_ in terms of the elastic constants *A*_1_, *A*_2_, *C* and *G*. MC calculations for the triad model which discretizes the continuum elasticity theory (3) were found to be in excellent agreement both with the perturbative (i.e., small *G*) and nonperturbative (for larger *G*) expressions of *C*_eff_. We note that despite being inaccessible experimentally, we were able to see the very high force behavior of the perturbative theory - namely the increase of *C*_eff_ from its low-force Moroz-Nelson behavior, towards its “naked” value of *C* in the triad MC calculations.

### B. oxDNA under moderate forces is described by the TWLC plus twist-bend coupling

To test whether our analytical results describe the coarse-grained behavior of a more realistic molecular model of DNA, we carried out MD simulations of oxDNA, a coarse-grained model describing DNA as two intertwined strings of rigid nucleotides [29]. *C*_eff_ for oxDNA1, a DNA model with symmetric grooves, was determined in previous work [34] and found to be in agreement with the (*G* = 0) Moroz-Nelson theory.

For the more realistic oxDNA2, which has the asymmetric grooves of real DNA and hence twist-bend coupling (*G* ≠ 0) [15], we found that *C*_eff_ is in excellent agreement with the nonperturbative theory Eq. (28) without any adjustable parameters, as the elastic constants were determined in previous work [15]. oxDNA2 appears to be precisely described by our nonperturbative theory for forces *f* ≪ *f*_0_ (recall that oxDNA undergoes internal structural transitions for forces of a few tens of pN, providing a more stringent contraint on force than the giant force scale *f*_0_). Put another way, the “TWLC plus *G*” is the “correct” low-force, long-fluctuation wavelength model for oxDNA2.

### C. Experimental data

We finally compare the analytical results with experimental magnetic tweezers data of Refs. [14, 22]. Figure 6 shows experimental data (symbols) together with plots of Eq. (28) for two sets of parameters *κ*_b_ and *κ*_t_ (lines). The latter are identical to the solid and dashed lines of Fig. 5, which are numerically precise descriptions of oxDNA1 and oxDNA2, respectively. Since the force fields in oxDNA were carefully tuned to reproduce several mechanical and thermodynamic properties of DNA [29], it is sensible to directly compare our nonperturbative theory to experimental data (Fig. 6).

**FIG. 6.**
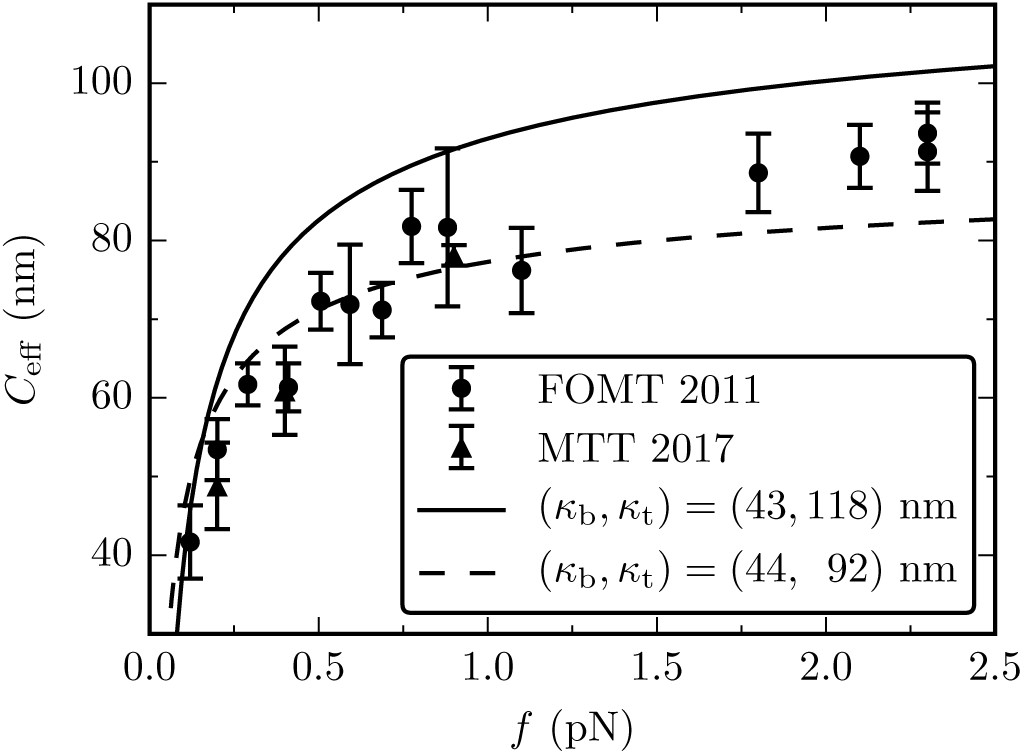
Comparison of the theory from Eq. (28) (lines) with Magnetic Tweezer experiments (symbols) for *C*_eff_ vs. force. The lines have the same parametrization of the dashed and solid lines of Fig. 5, which fit extremely well oxDNA and oxDNA2 data. Two sets of experiments are shown: the freely-orbiting magnetic tweezers [22] (circles) and magnetic torque tweezers [14] (triangles).

As reported in previous papers [14, 17] the experimental *C*_eff_ data are systematically lower than the prediction of the Moroz and Nelson theory, which precisely matches the oxDNA1 results (solid line in Fig. 6). The oxDNA2/nonperturbative theory (dashed curve) is closer to the experimental data, especially in the low force regime *f* < 1 pN. However, some systematic deviations are noticeable at higher forces, where theory appears to underestimate the experimental *C*_eff_. In addition, measurements at *f* = 15 pN (albeit for a slightly different assay) yield *C*_eff_ = 110 nm [35], well above the oxDNA2 value of *C*_eff_ = 92 nm.

We conclude by noting that oxDNA2 - which has the realistic features of groove asymmetry and G 6 = 0 - produces data in reasonable agreement with experiments. We have also shown that in the force range where we expect that coarse-graining of oxDNA2 should agree with our analytical results, it does. In that same force range(f < 10 pN), oxDNA2 and our analytical results show some systematic differences that suggest that physics beyond simple harmonic elasticity may be in play at intermediate forces (1 to 10 pN), generating torsional stiffening of DNA. This new ingredient might be sub-helix-scale structural transitions as discussed in [36], and highlighted in a recent study of thermally-driven DNA unwinding [28]. The next generation of coarse-grained DNA models likely will have to consider this kind of additional, internal degree of freedom to properly describe severe distortion of DNA by proteins, or similar situations where strong forces are applied at short length scales.

## ACKNOWLEDGMENTS

Discussions with M. Laleman and T. Sakaue are gratefully acknowledged. SN acknowledges financial support from the Research Funds Flanders (FWO Vlaanderen) grant VITO-FWO 11.59.71.7N, and ES from KU Leuven Grant No. IDO/12/08. JFM is grateful to the Francqui Foundation (Belgium) for financial support, and to the US NIH through Grants R01-GM105847, U54-CA193419 and U54-DK107980.

## Appendix A TWLC at strong stretching

We will first consider the simple case of the TWLC (*G* = 0), following closely the approach of Ref. [1]. At high forces, the molecule is strongly oriented along the force direction, which is chosen to be parallel to **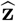**. It proves convenient to decompose the tangent vector as

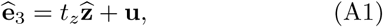

where the vector **u** is orthogonal to 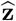, i.e. 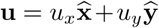. Using the identity 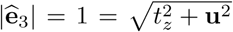 and expanding to lowest order in **u** we get

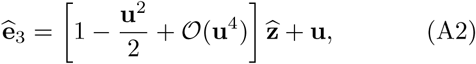

while its derivative is found to be

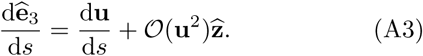

Combining this with Eq. (1), we find

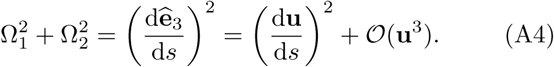

Introducing the Fourier transform 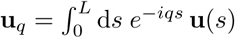, and neglecting higher-order terms, we write the bending and stretching contribution to the energy as follows

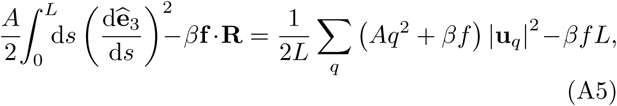

where we expressed the force as 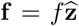, while the end-to-end vector 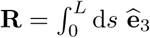 was approximated based on Eq. (A2).

The torque in Eq. (5) is coupled to the linking number, which is the sum of twist and writhe [Eqs. (7) and (8), respectively]. In the high-force limit, Eq. (8) becomes

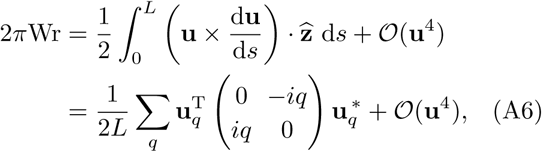

where we have rewritten the cross product as a matrix multiplication. Thus, the writhe couples the *x* and *y* components of the two-dimensional vector **u***_q_*.

Adding up all terms, and with the help of simple algebraic manipulations, we obtain the following energy for the TWLC to lowest order in **u**

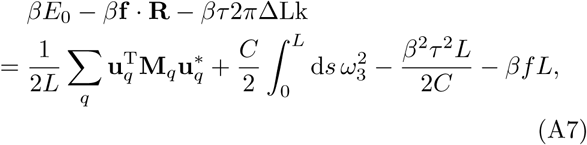

where we introduced the matrix

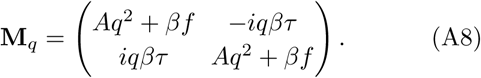

We have also introduced the shifted twist density

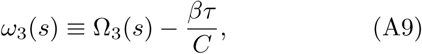

which allowed us to eliminate linear terms in Ω_3_. Thus, in the high-force limit, the TWLC under applied torque reduces, to lowest order, to a Gaussian model, where bending (**u***_q_*) and twist (*ω*_3_) are independent variables. The torque *τ* couples to the bending degrees of freedom through the off-diagonal terms of the matrix **M***_q_*. The eigenvalues of **M***_q_* are easily found to be

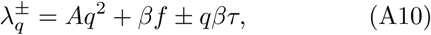

and the corresponding eigenvectors are 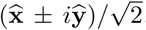. Writing Eq. (A7) on this basis allows us to calculate the partition function, from which the free energy is found to be

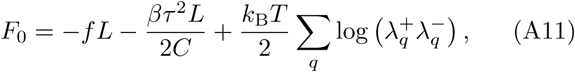

where we have neglected additive constants. Expanding to quadratic order in *τ*

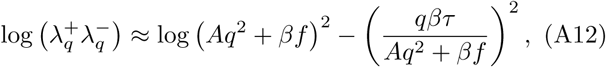

and replacing the sum over momenta with an integral ∑*_q_* → (*L*/2*π*) ∫ d*q*, we obtain

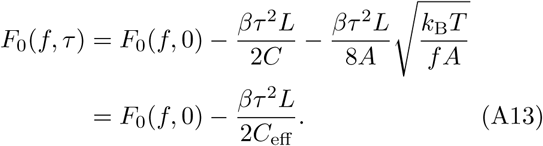

Combining the two last terms in the right-hand side, one obtains the Moroz and Nelson relation [Eq. (10)]. *F*_0_(*f, τ* = 0) is the zero-torque free energy, and is obtained by integrating the first term at the right-hand side of Eq. (A12). Although the integral is divergent, it can be regularized by introducing a momentum cutoff Λ ≈ 2*π/a*, where *a* = 0.34 nm is the separation between neighboring base pairs. As it turns out, however, this cutoff does not affect any force-dependent terms, and one has

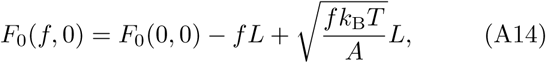

where *F*_0_(0, 0) is cutoff-dependent. Interestingly, from Eq. (5) one finds to lowest order in *τ*

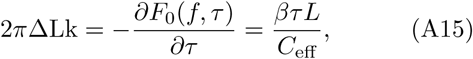

which quantifies the induced over- or undertwisting upon the application of a torque. Using this expression, one obtains the force-extension relation at fixed linking number [4, 21]

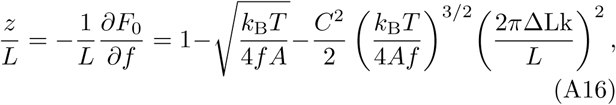

which shows a characteristic parabolic profile for the extension of an over- or undertwisted, stretched molecule.

## Appendix B Ω2 at strong stretching

The thermal average in Eq. (12) contains Ω_2_, which needs to be expressed in terms of **u***_q_* and *ω*_3_, the degrees of freedom of the system [Eq. (A7)]. For this purpose, we use the relation

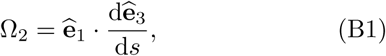

which can be easily obtained from Eq. (1). In the high-force limit, where the tangent ****ê****_3_ points predominantly along the force direction, 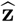, one has

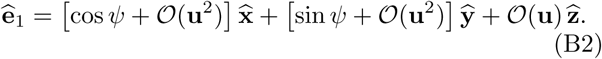

Here we have introduced the twist angle

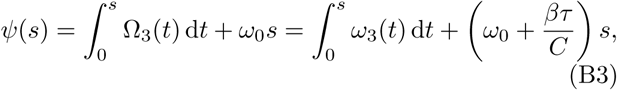

and used Eq. (A9) to express it in terms of the variable *ω*_3_. Equation (B2) can be obtained by considering an arbitrary rotation that maps a fixed lab frame triad, e.g. 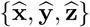, onto the material frame triad {****ê****_1_, ****ê****_2_, ****ê****_3_} at position *s*, requiring that ****ê****_3_ remains predominantly oriented along the force direction, as in Eq. (A1). Combining Eqs. (B1) and (B2), it follows that

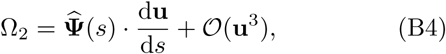

where we have defined the unit vector

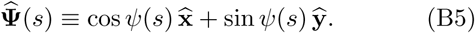

Therefore, in the high-force limit, Ω_2_ can be written as a scalar product between a unit vector 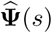, depending exclusively on twist variables, and a vector du/d*s*, involving only the bending degrees of freedom. Finally, from Eq. (B4) it follows that

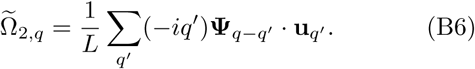

The remainder of the calculation, presented below, will be based upon Eqs. (B4) and (B6).

## Appendix C Details of the perturbative calculation

To calculate the average appearing in Eq. (12), it first needs to be rewritten as a function of the integration variable *ω*_3_ [see Eq. (A9)]. This can be performed as follows

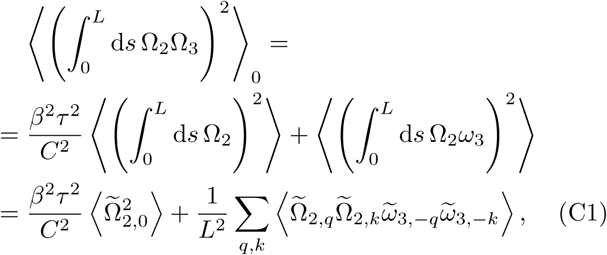

where 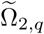 and 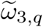 denote the Fourier components of Ω_2_ and *ω*_3_, respectively. Note that we have neglected a linear term in *ω*_3_, which vanishes due to the symmetry *ω*_3_ ↔ *-ω*_3_. Moreover, in order to simplify the notation, we have dropped the subscript from all averages ⟨.⟩_0_, which will be always calculated within the TWLC model, i.e. for *G* = 0.

Before proceeding to the calculation of Eq. (C1), it will prove useful to first present some properties. In particular, we are going to use the following expressions, obtained from the correlation functions in the TWLC model [Eq. (A7)]

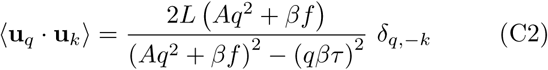

and

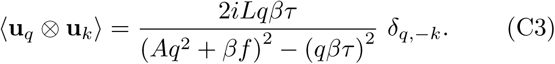

For convenience, we have introduced the shorthand notation

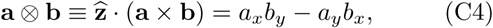

which is antisymmetric with respect to the interchange of **a** and **b**. From Eq. (C3) it follows that ⟨**u***_q_* ⊗ **u***_k_* ⟩= 0, when *τ* = 0 (in this case the matrix **M***_q_* is diagonal, hence the cross-correlations 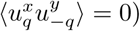. Moreover, for *τ* = 0 and *q* = −*k*, Eq. (C2) reduces to:

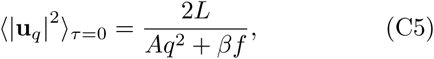

which can be easily obtained from equipartition [12]. We are also going to use the following symmetries

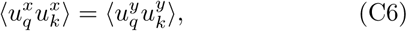

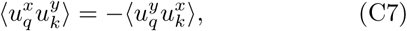

which allow us to rearrange scalar products as follows

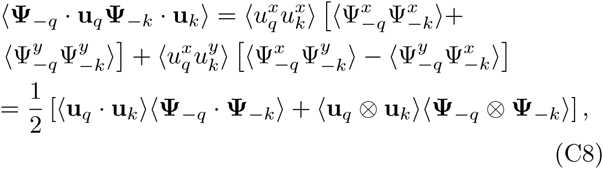

where we have used the fact that the bending (**u**) and twisting (**Ψ**) degrees of freedom are independent, within the TWLC. We are now ready to proceed to the calculation of Eq. (C1). We will need to evaluate two distinct terms, which will be treated separately.

### 1. First term in Eq. (C1)

The first term in Eq. (C1) already contains a factor of order 𝒪(*τ* ^2^), which means that up to quadratic order in *τ* it is sufficient to evaluate the corresponding average for *τ* = 0

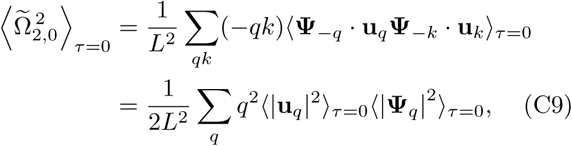

where we have used Eqs. (B6) and (C8), together with the property ⟨ **u***_q_* ⨂ **u***_k_* ⟩ _*τ*=0_ = 0 [see Eq. (C3)]. Next, we need to calculate the following quantity

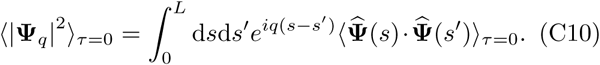

From Eqs. (B3) and (B5) one finds

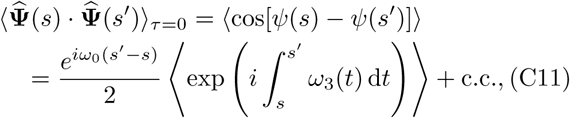

where c.c. denotes the complex conjugate. To proceed, we perform a Fourier transform of the exponent

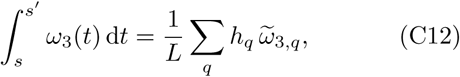

where we have introduced the complex variable

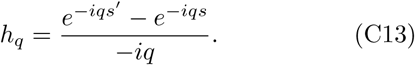

Performing Gaussian integration in 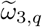, one finds

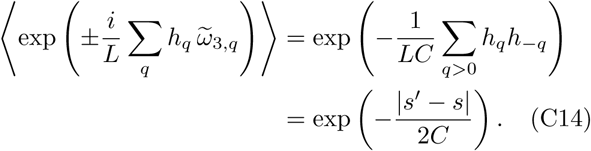

As expected, the decay of the twist correlation function is governed by 2*C*, i.e. the twist persistence length in the TWLC. Combining Eqs. (C11), (C12) and (C14), we obtain

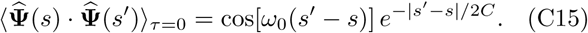

Inserting this in Eq. (C10) yields

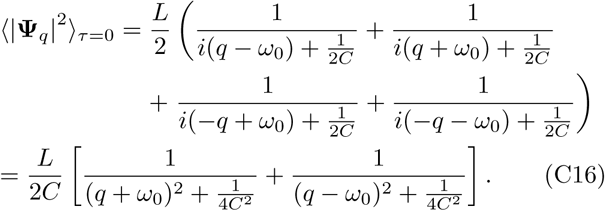

Finally, combining Eqs. (C9), (C5) and (C16) we find

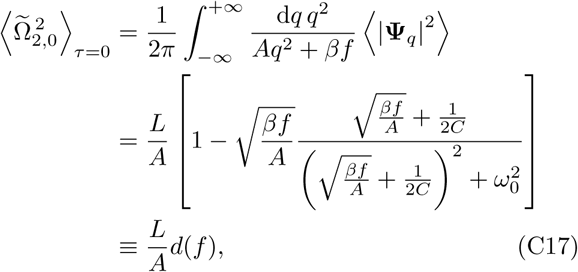

where we introduced a force-dependent scale factor *d*(*f*). Note that 0 ≤ *d*(*f*) ≤ 1, with *d*(*f*) → 1 at small forces and *d*(*f*) → 0 at high forces. Commonly accepted estimates of the DNA elastic constants put them in the viccinity of *C* = 100 nm and *A* = 50 nm, while the applied forces in typical experiments are in the range 0.1 pN ≲ *f* ≲10 pN. Recalling that room temperature corresponds to *k*_B_*T* ≈ 4 pN·nm, it follows that *βf/A* is at least one order of magnitude larger than 1/4*C*^2^. This allows for the following simplification

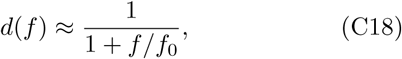

neglecting higher-order terms in 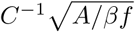. We have also introduced a characteristic force

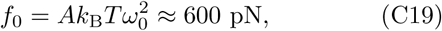

whose value greatly exceeds those at which the double helix breaks. Thus, for the force range of interest, we may set *d*(*f*) = 1 in Eq. (C17). Summarizing, this first term of Eq. (C1) provides the following contribution to the free energy

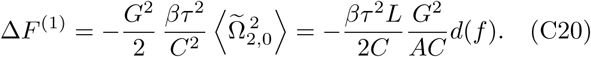

As a final remark, we note that we could have obtained Eq. (C18) using the approximation

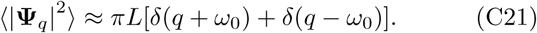

Formally, this corresponds to taking the limit *C* → ∞ in Eq. (C16), i.e. approximating the Lorentzian distributions 1/[2*C*(*q* ± *ω*_0_)^2^ +1/2*C*] with delta functions. This is a valid approximation as long as *ω*_0_ ≫ 1/2*C*, making the Lorentzians sharply-peaked at large momenta *q* = ± *ω*_0_, where the integrand in Eq. (C17) varies slowly.

### 2. Second term in Eq. (C1)

Using the same decomposition as in Eq. (C8), the second term in Eq. (C1) can be written as

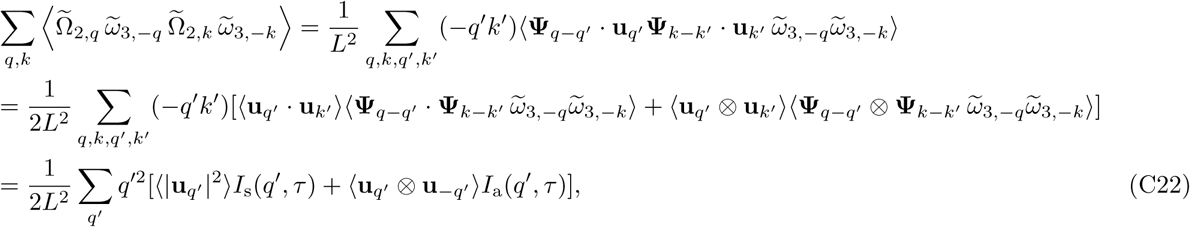

where we used the fact that the **u** correlators are diagonal in momentum space, hence *k* ′ = – *q*′ [see Eqs. (C2) and (C3)]. We have also introduced the symmetric

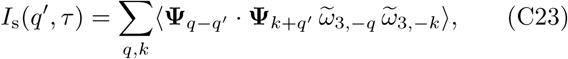

and antisymmetric products

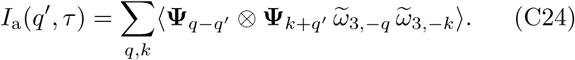

In what follows, we are going to compute the contribution of *I*_s_ and *I*_a_ to the free energy separately.

#### a. Symmetric products

For the evaluation of Eq. (C23), we will first focus on the average inside the summation, which may be written in the following way

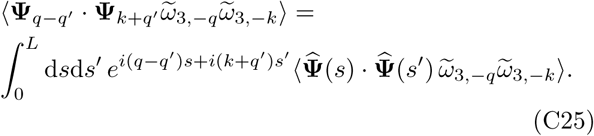

we may now use Eqs. (C11)-(C14) so as to obtain

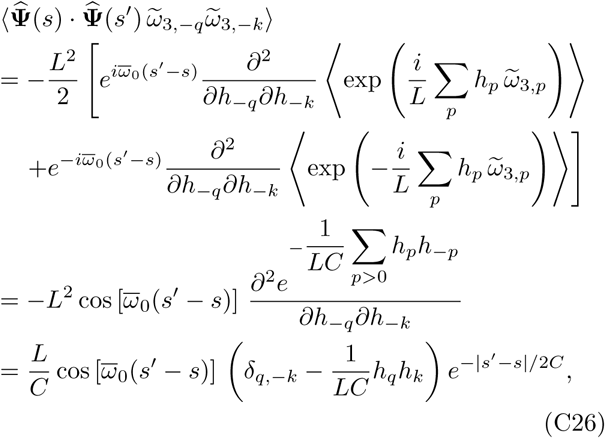

where we have introduced the shifted intrinsic twist

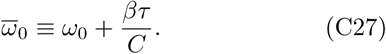

Differently from the calculation of 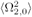 in Eq. (C9), we can no longer ignore the torque dependence of *ψ* [see Eq. (B3)]. Plugging Eq. (C26) back in Eq. (C25), integrating in *s* and *s*′ and summing over *q* and *k*, we obtain

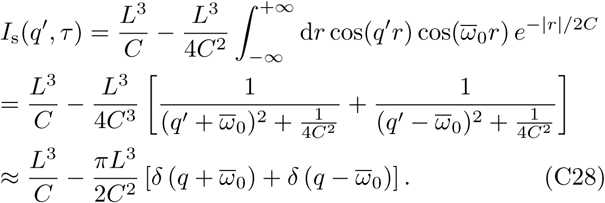

Throughout the calculation we introduced the variable *r* ≪ *s*′ – *s* in the double integral. Similar to Eq. (C21), we also approximated the two Lorentzians with delta functions. Note that *I*_s_ depends on the torque *τ* through *ω*_0_, as indicated by Eq. (C27). Combining Eqs. (C2) and (C28), one finds

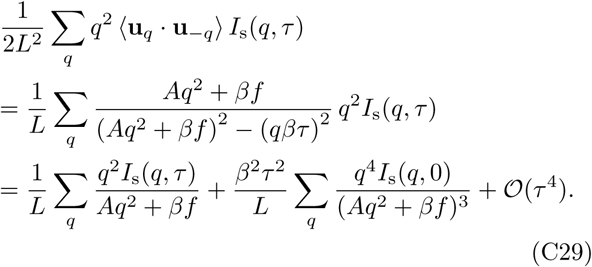

we are interested in terms proportional to *τ* ^2^. There are two such contributions, the first one being

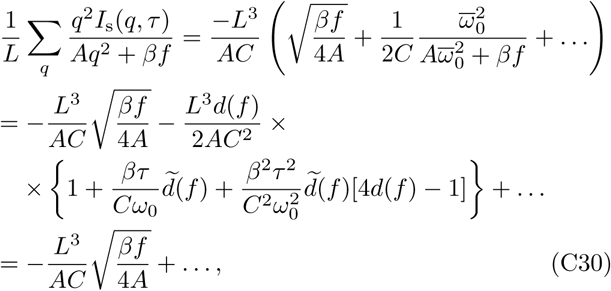

where we defined 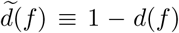, with *d*(*f*) the scale factor given in Eq. (C18), and where the dots indicate omitted terms, which do not significantly contribute to the result. These terms are either independent of the torque and force, or are of higher order than *τ* ^2^. We note that 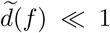, i.e. it is negligibly small for the experimentally-accessible forces *f* ≪ *f*_0_ 600 pN. The only surviving term i<u>n</u> Eq. (C30) is independent of *τ* and proportional to 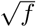, hence contributing to the force-extension response. The remaining term to evaluate in Eq. (C29) is

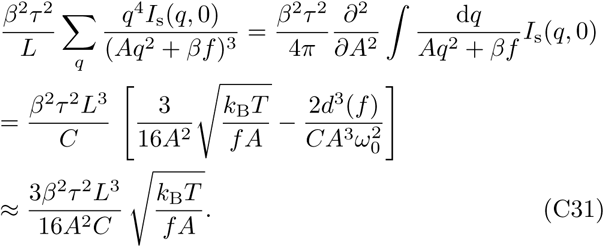

Note that terms containing 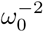 are always multiplied by *A*^−2^ or *C*^−2^, hence forming dimensionless constants. Typical values for the case of DNA are (*Aω*_0_)^−2^ ≈ 10^−4^ and (*Cω*_0_)^−2^ ≈ 3 × 10^−5^, which provide negligible contributions to the free energy, compared to other terms of the same order in *τ*. Therefore, the term proportional to *d*^3^ in Eq. (C31) can be safely neglected. Combining Eqs. (C29)-(C31), we find that the relevant contribution of the symmetric term in Eq. (C22) to the free energy is

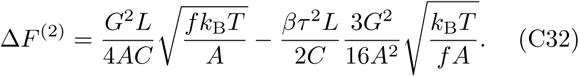

#### b. Antisymmetric products

The final part of the derivation is devoted to the calculation of the antisymmetric product in Eq. (C22). We start by expanding Eq. (C3) as follows

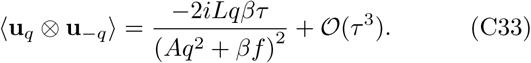

The calculation of the twist correlator is performed in a similar fashion as above, which yields

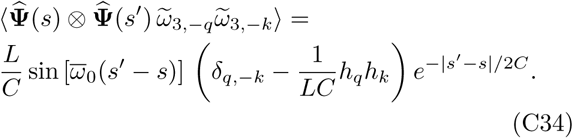

We may take the Fourier transform of this expression, and plug it back into Eq. (C24), so as to obtain

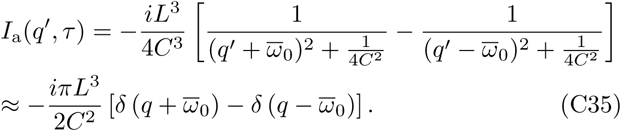

Finally, plugging Eqs. (C33) and (C35) into the second term of Eq. (C22), transforming the sum into an integral and performing the remaining integration, we find

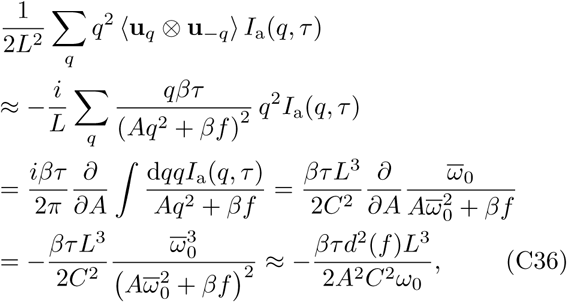

where we have omitted terms, which are either higher order in *τ*, or negligibly small compared to other terms of the same order [recall (*Cω*_0_)^−2^ 3 10^−5^]. Summarizing, the contribution of the antisymmetric product to the free energy is

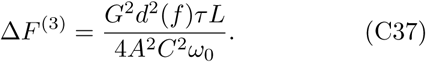

### 3. Collecting the results

Throughout the derivation we found three distinct contributions to the free energy, coming from Eqs. (C20), (C32) and (C37). Adding these to Eq. (A13), i.e. the free energy of the TWLC, we find

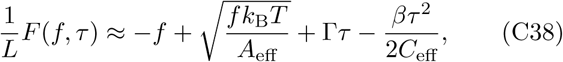

where we have omitted both terms independent of *f* and *τ* and higher-order corrections in *τ* and *G*. We have also introduced the effective bending stiffness

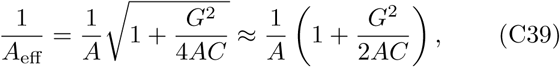

together with the proportionality constant

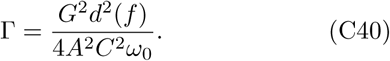

Finally, we reach the following expression for the effective torsional stiffness

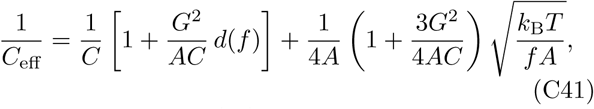

corresponding to Eq. (16) of the main text, and the central result of this work.

## Appendix D Intrinsic bending

The analysis above is based on description of the ground-state configuration of DNA relative to a straight molecular axis [**Ω** = **0** in Eq. (1)], i.e., for a molecular axis which is straight in the ground state. However, one can also choose coordinates where the ground state of the double helix is a helix while still respecting the symmetry of the elastic model. In fact, this is a rather natural outcome for most choices of DNA deformation which are based on molecular modeling, where coordinates are usually chosen relative to the orientations of the base pairs (e.g., using the vector connecting the junctions of the bases to the sugar-phosphate backbone as a reference), due to the groove asymmetry of DNA. Most relevant here, our previous determination of the elastic constants of oxDNA2 [15] analyzed deformations relative to a helical coordinate system. We now show how to transform the elastic constants in such a helical coordinate system to the straight-line coordinates relevant to our calculations.

Intrinsic bending consistent with groove asymmetry, usually reported in the DNA literature as a nonzero value of the average roll [37], can be described using the following modification of Eq. (1)

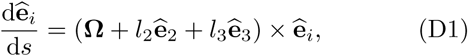

where *l*_2_ and *l*_3_ correspond to the intrinsic bending and twisting densities, respectively, with *l*_2_ ≪ *l*_3_. A nonzero *l*_1_ is incompatible with the symmetry of the double helix.

Solving Eq. (D1) for **Ω** = **0**, one finds that the ground-state configuration is a helix, with a linking number equal to Lk_0_ = *ω*_0_*L*/2*π*, where

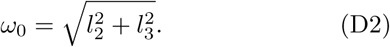

Furthermore, from the solution of Eq. (D1), it follows that the rotation matrix transforming the helical ground state of Eq. (D1) to the straight one of Eq. (1) is

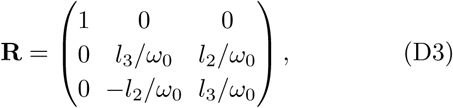

expressed on the body frame {****ê****_1_, ****ê****_2_, ****ê****_3_} of the former.

The total elastic free energy should not depend on the coordinate system used to describe it, so the energy in the helical coordinates (*l*_2_, *l*_3_ ≠ 0) should equal that found in non-helical coordinates (*l*_2_ = 0 and *l*_3_ = *ω*_0_) The deformations in the “straight” model are given by **Ω ′** = **R**^T^**Ω**, where **Ω** are the deformation parameters of the helical model. From the condition that the integrand of Eq. (2) has to remain invariant under this transformation, one obtains the following relations mapping the elastic constants from the helical coordinates to the straight ones:

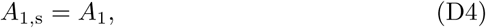

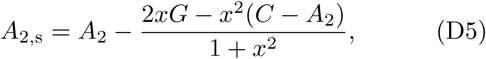

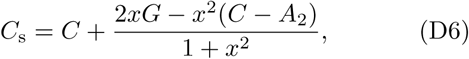

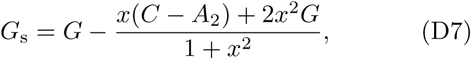

where *x* ≡ *l*_2_*/l*_3_ ≪ 1, and where the subscript s indicates the “straight” frame result. The transformation (D3) mixes *A*_2_, *G* and *C*, changing their values, but conserves the symmetry of the elastic constant matrix. These formulae allow one to measure elastic constants using arbitrarily chosen helical reference coordinates, and then convert them to elastic constants suitable for using strains defined relative to a straight-line ground state.

For unconstrained (zero force and torque) oxDNA2 simulations, we measured reference helix parameters *l*_2_ = 0.1349 nm^−1^ and *l*_3_ = 1.774 nm^−1^, giving *ω*_0_ = 1.779 nm^−1^ and *x* = 0.076. Elastic constants reported in Ref. [15] (*A*_1_ = 85 nm, *A*_2_ = 39 nm, *C* = 105 nm, and *G* = 30 nm) were measured in reference to helical coordinates; for use in our analytical theory we transform them to the straight coordinates using (D4)-(D7) to obtain the oxDNA2 values in Table 1.

